# Spatially resolved analysis of longitudinal fruit growth in persimmon (*Diospyros kaki*) via three-dimensional phenotyping

**DOI:** 10.1101/2025.07.17.665444

**Authors:** Akane Kusumi, Soichiro Nishiyama, Hisayo Yamane, Ryutaro Tao

## Abstract

Understanding fruit development patterns is essential, as they are directly linked to fruit size and shape, which ultimately determine final yield and quality. While conventional models describe fruit development based on growth parameters for the whole fruit, it remains unclear how tissue growth at different fruit portions is coordinated to promote overall fruit development. Therefore, in this study, we investigated the relationship between differences in developmental rates among fruit portions and fruit morphology in persimmon (*Diospyros kaki*), a species known for its high morphological diversity. Starting two weeks after blooming (WAB), surface landmarks were routinely drawn along the calyx-apex line on fruits of four persimmon cultivars with diverse shapes, and the landmark displacement was quantified every two weeks using 3D phenotyping. Consistent with previous studies, all cultivars exhibited active development in the regions near the calyx, with the greatest difference in growth rates between the calyx and apex occurring at 6–8 WAB. However, the extent of these spatial differences varied among cultivars. In elongated fruits, a gradual growth gradient was observed from the calyx toward the apex, with relatively high growth rates even near the apex. In contrast, in flattened fruits, growth was highly concentrated near the calyx, while development in other regions remained minimal. The findings of this study provide a foundation for future research on fruit shape regulation and the elucidation of physiological disorders in persimmon.

**Highlights:**

- Spatially resolved 3D phenotyping quantified growth dynamics in persimmon fruit
- Growth rates varied along the fruit axis, with active growth near the calyx in all tested cultivars
- Growth gradients were linked to final fruit shape

## 1. Introduction

Fruit development is a fundamental research area in horticulture, as deviations in growth influence fruit size and shape, thereby ultimately impacting both yield and quality. The development of fleshy fruit, which accounts for majority of commercial fruit production, is generally driven by cell division in the early stages, followed by cell expansion (Gillaspy et al., 1993). Several key genes that regulate the rate and orientation of cell division and expansion have been identified as determinants of final fruit size and shape (van der Knaap et al., 2014). Through the coordinated regulation of these cellular processes, fruit development, as represented by changes in fruit size and weight, typically follows either a single sigmoid growth curve or a double sigmoid growth curve. The single sigmoid growth pattern, observed in pome fruits such as apple and pear (Tukey and Young, 1942; Crane, 1964), is characterized by an initial phase of rapid cell division, followed by cell expansion at later stages. In contrast, the double sigmoid growth curve, seen in fruits such as blueberry (Young, 1952) and sweet cherry (Tukey and Young, 1939), includes a distinct growth suspension phase during mid-development. This temporary halt in growth is attributed to the hardening of endocarp and embryo development, leading to competition for nutrients between the embryo and surrounding fruit tissues, ultimately suppressing fruit expansion (Yamaguchi et al., 2002). Understanding these fruit development patterns has long been essential for optimizing orchard management and improving crop production.

Although the fruit growth curves primarily describe overall fruit development well, they do not explicitly examine how the growth dynamics of individual tissues within the fruit contribute to its overall development. Few studies have investigated the relationship between the spatial localization of growth within the fruit and the final fruit morphology. Eldridge et al. (2016) examined tissue-specific developmental dynamics that contribute to the different fruit shape by analyzing cell division patterns and subsequent growth in Brassicaceae species. In the cylindrical *Arabidopsis thaliana* silique, tissue growth rates and orientations were relatively uniform along the proximal-distal axis, while they were non-uniform in the heart-shaped *Capsella rubella* silicle. A subsequent study revealed that *SHOOTMERISTEMLESS* (*STM*) was highly expressed at the bulge of the *Capsella* fruit apex, contributing to the maintenance of local cell division (Hu et al., 2025). Such insights into tissue-specific growth regulation have been facilitated by the ease of genetic transformation in model plants. However, in horticultural crops especially in fruit trees, research on tissue-level developmental dynamics remains limited, highlighting the need for further studies to elucidate the mechanisms governing fruit shape formation.

The cultivated oriental persimmon (*Diospyros kaki*) is a temperate deciduous fruit tree crop characterized by distinctive genetic diversity in fruit growth and morphology (Kusumi et al., 2024). The fruit development of persimmon follows a double sigmoid growth curve, with an initial stage I of rapid fruit enlargement after flowering, followed by stage II of growth suspension, which typically occurs from August to September. Thereafter, the fruit enters stage III, the final stage of development (Nakano et al., 1998).

One of the distinctive features of persimmon fruit development is the vigorous growth observed at the calyx-end. Fujimura (1935) drew circle landmarks on the base, apex, and middle of ‘Fuyu’ fruit in early July and measured their diameters every two weeks. He found that these circle landmarks shifted toward the fruit apex over time, with those near the base exhibiting a rapid increase in both lateral and vertical diameters. Similarly, Yamamura (1984), in his investigation into the relationship between the susceptibility of ‘Saijo’ fruit to black stain on its skin and its growth characteristics, drew parallel lines at 2 mm intervals on ‘Saijo’ and ‘Fuyu’ fruit approximately 25 days after full bloom. It was reported that the growth of the basal region was more pronounced compared to that of the apical region. These previous studies indicate that the basal region of persimmon fruit undergoes particularly active growth. However, none of these studies have explicitly examined the relationship between growth dynamics and fruit morphology across cultivars with diverse shapes found in different genotypes of persimmon. Therefore, a comparative analysis among cultivars is expected to provide novel insights into the factors influencing fruit shape development.

This study aims to visualize differences in developmental rates among fruit regions and to quantify the effects of localized tissue growth dynamics on persimmon fruit development. Such an approach will serve as a foundation for research on fruit size and shape determination and may ultimately contribute to the control of fruit size or those with desired shapes. Furthermore, this study is expected to aid in elucidating the causes and predicting the occurrence of physiological disorders common in persimmon, such as calyx-end cracking (*hetasuki*) and apical-end splitting (Yamada et al., 1988), as well as fruit malformations.

## 2. Materials and Methods

### 2.1 Plant materials and fruit surface marking

Fruits from four persimmon cultivars, ‘Saijo’, ‘Yamato’, ‘Hiragaki’, and ‘Tamopan’, planted in Kyoto Farmstead of the Experimental Farm of Kyoto University were investigated during the 2022 season. The experiment began at 2 weeks after blooming (WAB) for ‘Saijo’, ‘Yamato’, and ‘Hiragaki’, while it started at 6 WAB for ‘Tamopan’. Surface landmarks were drawn along three lines connecting the calyx and apex on one-quarter of the fruit, corresponding to the portion derived from a single carpel, using an ink marker. Using a paper or string guide pre-marked at 3.4 mm intervals, landmarks were drawn at each 3.4 mm interval along the calyx-apex line, starting from the fruit apex as the origin and extending toward the calyx until the track overlapped (Fig. 1A). To account for the increase in fruit size during development, landmarking was repeatedly applied to new fruits at regular intervals, with an increasing number of marks per line. The schematic of the marking and sampling schedule is shown in Fig. 1B. Four to five marked fruits were sampled every two weeks. To ensure complete fruit shape capture, the calyx was removed from the sampled fruits before 3D scanning.

**Fig. 1.**
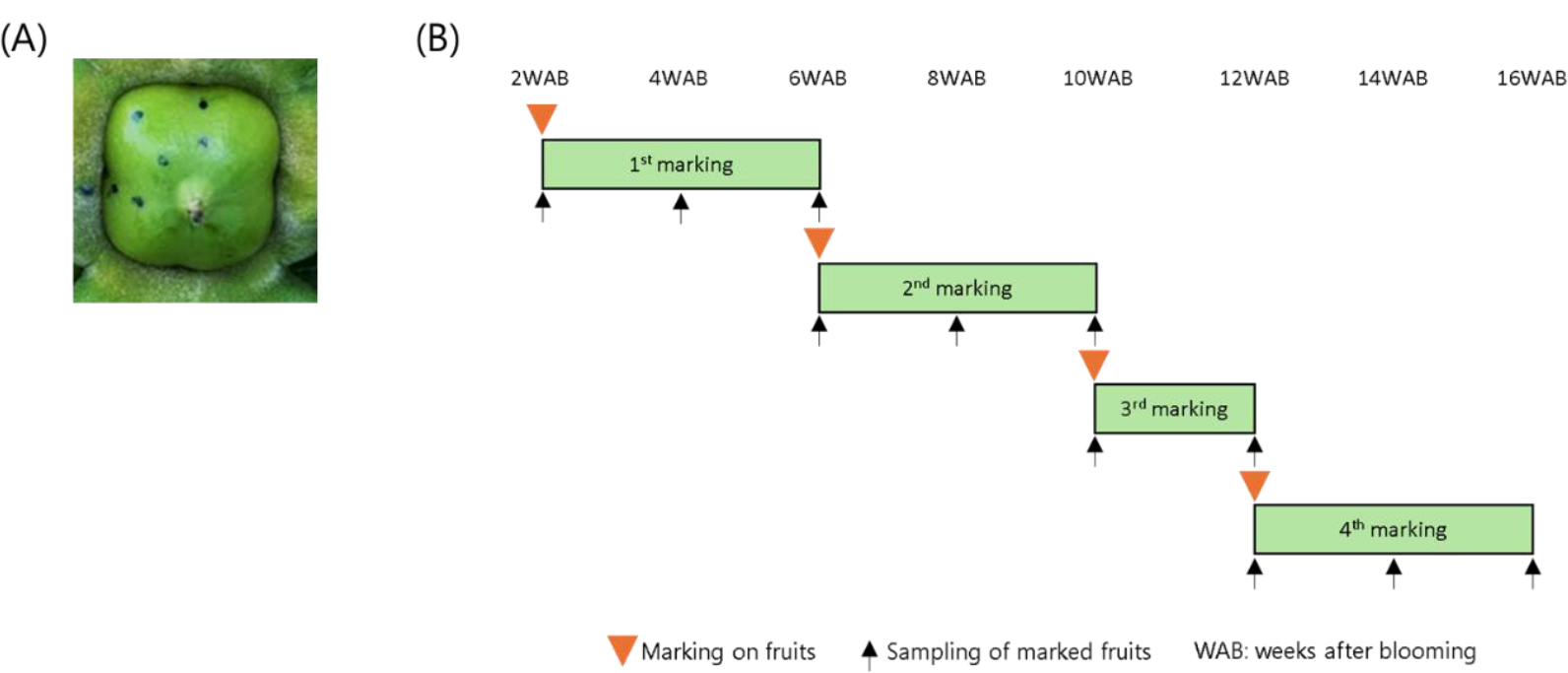
Spatial fruit growth analysis by surface landmarking in this study. (A) Example of surface landmarking on *‘*Saijo*’* fruit at 2 weeks after blooming (WAB). Three dotted lines were drawn at 3.4 mm intervals with the fruit apex as an origin until it overlapped the calyx. (B) The base timeline for routine fruit landmarking and sampling. It was adjusted according to each cultivar.

### 2.2 Reconstruction of 3D models and measurement of the landmarks

A 3D Scanner Coordinate Measuring Machine (VL-500; KEYENCE, Osaka, Japan) was used to generate complete 3D models of the sampled persimmon fruit. We previously reported that the mean absolute distance error of 0.116±0.0318 mm between physical and digital point-to-point measurements (n=10) and a scan-to-scan repeatability of 0.0173 mm (n=3) for fruit phenotyping, confirming sub-millimeter accuracy (Kusumi et al, 2024). Upon the measurement, the fruit apex was set as the origin (mark zero), and subsequent marks were numbered sequentially, starting from the one closest to the apex.

The distance *d*_n_ between the n^th^ and (n-1)^th^ landmarks from the fruit apex (n ≥ 1, with the fruit apex as the 0^th^ mark) along a dotted line was calculated in 3D space for ‘Saijo’, ‘Yamato’, and ‘Hiragaki’ using the software accompanying the VL-500 (Fig. 2). For ‘Tamopan’, which exhibits a more complex fruit shape, sections of the 3D model were extracted along the dotted line, and the distance *d* between adjacent landmarks on the fruit contour was measured in 2D space. This adjustment was necessary because the presence of horizontal grooves led to discrepancies between the direct 3D-measured distances and the actual distances along the fruit contour.

**Fig. 2.**
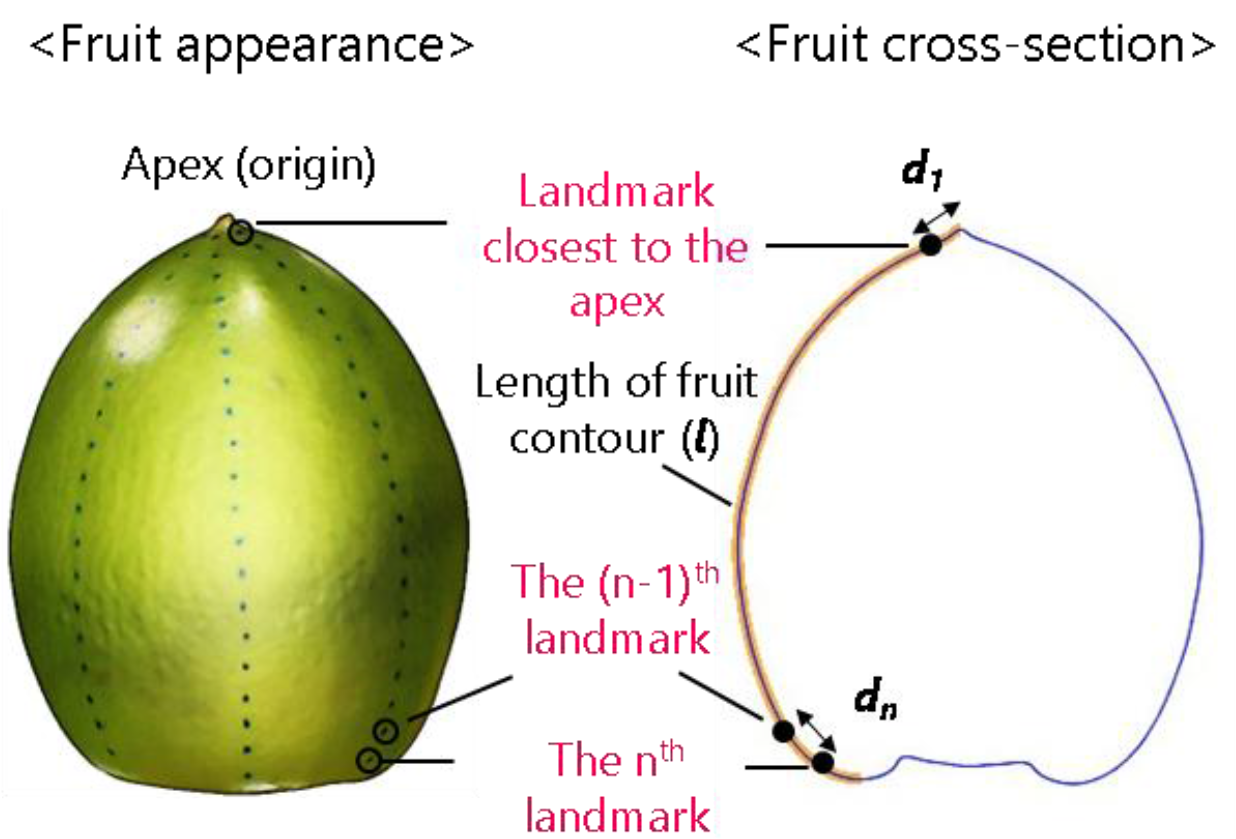
Measurement of the surface landmarks to model spatial dynamics of fruit development in persimmon. Distance *d* between two adjacent landmarks and *l* were used to calculate the relative position of the landmark.

Additionally, the relative position of each landmark along the fruit contour was determined. First, the contour length (*l*) between the fruit apex and the fruit shoulder in a cross-section obtained along the dotted line was measured for all cultivars (Fig. 2). The relative position *p*_n_ of the n^th^ landmark on the fruit contour was defined as:

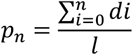

where the landmarks closer to the fruit apex have values approaching 0, and those nearer the fruit calyx have values approaching 1.

### 2.3 Analysis of fruit growth

Based on the information of *d*_*n*_ and *p*_*n*_, two aspects of fruit growth were analyzed. The first aspect involved examining changes in the relative positions of the landmarks closest to the calyx and the apex. These changes were compared with those from the previous sampling point by subtracting the relative position of the landmark at present from the one at the previous time. Therefore, if the landmark moved to the apex end, its change was regarded positive.

The second aspect focused on changes in the distances between adjacent marks across different fruit regions. Since data collection was conducted through destructive sampling, direct comparisons between replicating fruits at the same sampling time and between consecutive sampling periods were challenging due to size and shape variability. To account for these variations, surface landmarks were categorized into three portions, namely distal (apex-end), middle, and proximal (calyx-end). This categorization helped to normalize measurement variability across fruits by averaging differences across similarly defined regions. Additionally, to minimize these errors, outliers were excluded from the whole *d*_n_ data at each sampling point using the interquartile range (IQR) method. Outliers were defined as values smaller than the first quartile minus 1.5 times the interquartile range or larger than the third quartile plus 1.5 times the interquartile range. After excluding these outliers, the remaining *d*_n_ data along the marked line were categorized into three portions—distal, middle, and proximal—and the average *d*_n_ was calculated for each portion.

## 3. Results

### 3.1 Analysis of fruit growth curves using 3D phenotyping

First, we analyzed fruit growth patterns of four cultivars to visualize their general growth characteristics based on 3D modeling. The seasonal changes in fruit volume, height and width were shown in Fig. 3. The growth curve of persimmon fruit is generally represented by a double sigmoid pattern (Nakano et al., 1998). The increasing pattern of fruit volume followed this double sigmoid pattern. ‘Saijo’, ‘Hiragaki’, and ‘Tamopan’ clearly indicated the presence of stalled growth (stage II) at around 12 WAB, while ‘Yamato’ did not show the clear double sigmoid growth curve (Fig. 3A). Horizontal and vertical fruit growth, represented by the increase of height and width respectively, also followed similar double sigmoid patterns (Fig. 3B and 3C). Fruit growth halted at around 12 WAB and this was more apparent in ‘Hiragaki’, ‘Tamopan’ and ‘Saijo’. This indicated that coordinated and largely synchronous growth along both longitudinal and lateral axes underlies an overall increase in fruit volume.

**Fig.3.**
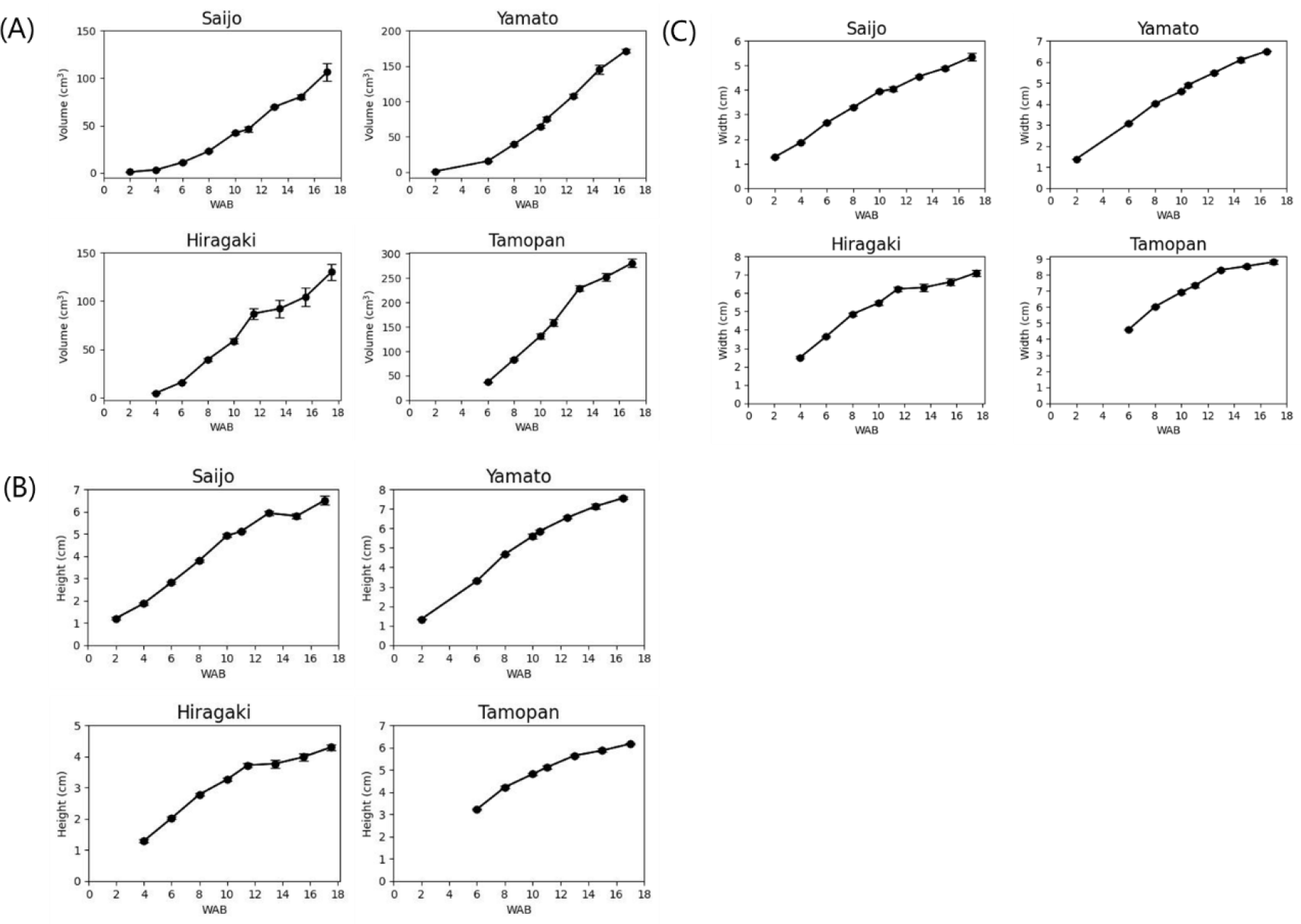
Fruit growth patterns of four cultivars in this experiment. Three parameters were measured, (A) fruit volume, (B) fruit height, and (C) fruit width. WAB stands for weeks after blooming. The values represent the average of more than three fruits, and the error bars indicate the standard error.

### 3.2 Variation in fruit growth at the proximal and distal ends

Fruit surface landmarks remained clearly visible even two weeks after they were applied. Reconstructed 3D models of the sampled fruits are presented in Fig. 4. A visual comparison between fruits immediately after marking at 6 WAB and those observed two weeks later revealed that the landmark closest to the proximal end of the fruit shifted away from the fruit shoulder and moved toward the distal end. In contrast, during the later stages of fruit development, such as around 13 WAB, the displacement of surface landmarks became less pronounced.

**Fig. 4.**
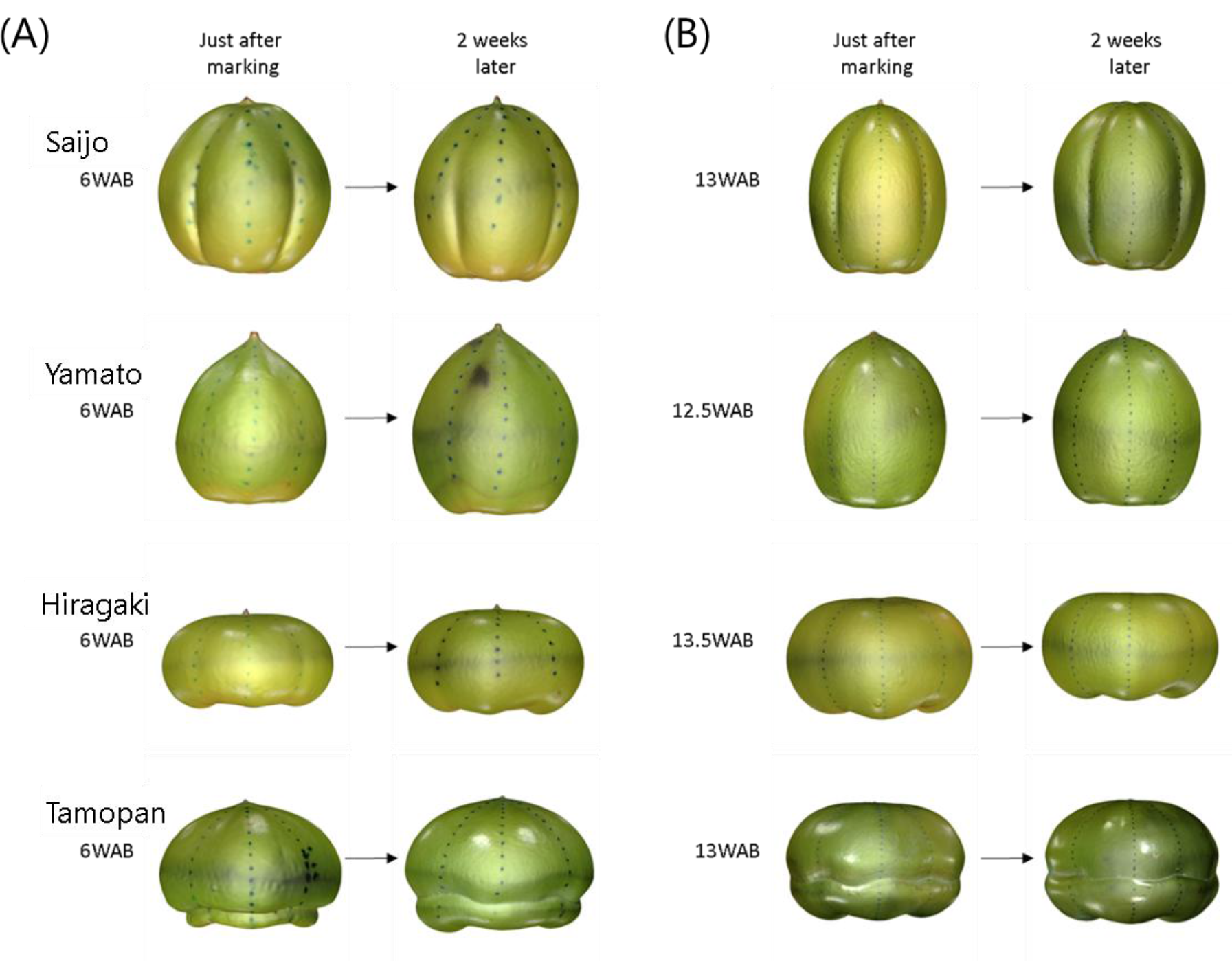
Reconstructed 3D models of marked fruits representing the fruit growth in 2 weeks. (A) Comparisons between the fruits just after being marked at 6 weeks after blooming (WAB) and the ones two weeks later. (B) Comparisons between the fruits just after the final landmarking and the ones two weeks later.

Temporal changes in the relative positions of the landmarks closest to the apex and calyx are shown in Fig. 5A. In the initial stages of fruit development, the landmark on the distal end appeared to shift slightly more than that on the proximal end, as observed in ‘Saijo’ and ‘Yamato.’ However, after 4 WAB, when cell expansion became more pronounced, the positional change of the landmark on the proximal end exceeded that of the distal end in all cultivars, reflecting the vigorous tissue development near the calyx. Growth on the proximal end peaked at approximately 6–8 WAB, whereas growth on the distal end reached its peak earlier. Thereafter, growth on both the apex and calyx sides gradually declined, approaching zero around 13 WAB. Later, in early September (approximately 16 WAB), a slight increase was observed in all cultivars except ‘Yamato.’ The tissue developmental dynamics represented by this temporal pattern of positional changes in surface landmarks closely resembled the growth curves for fruit volume and height. Specifically, tissue development, particularly near the distal end, exhibited a high growth rate in the early stages of fruit development, slowed during the second stage, and then showed a slight increase again in the third stage. Notably, the extent of growth changes between the calyx and apex sides varied among cultivars (Fig. 5B). In all cultivars, the greatest differences were observed at 6–8 WAB, with the largest value recorded in ‘Hiragaki’ (0.127), followed by ‘Tamopan’ (0.072), ‘Saijo’ (0.047), and ‘Yamato’ (0.038).

**Fig. 5.**
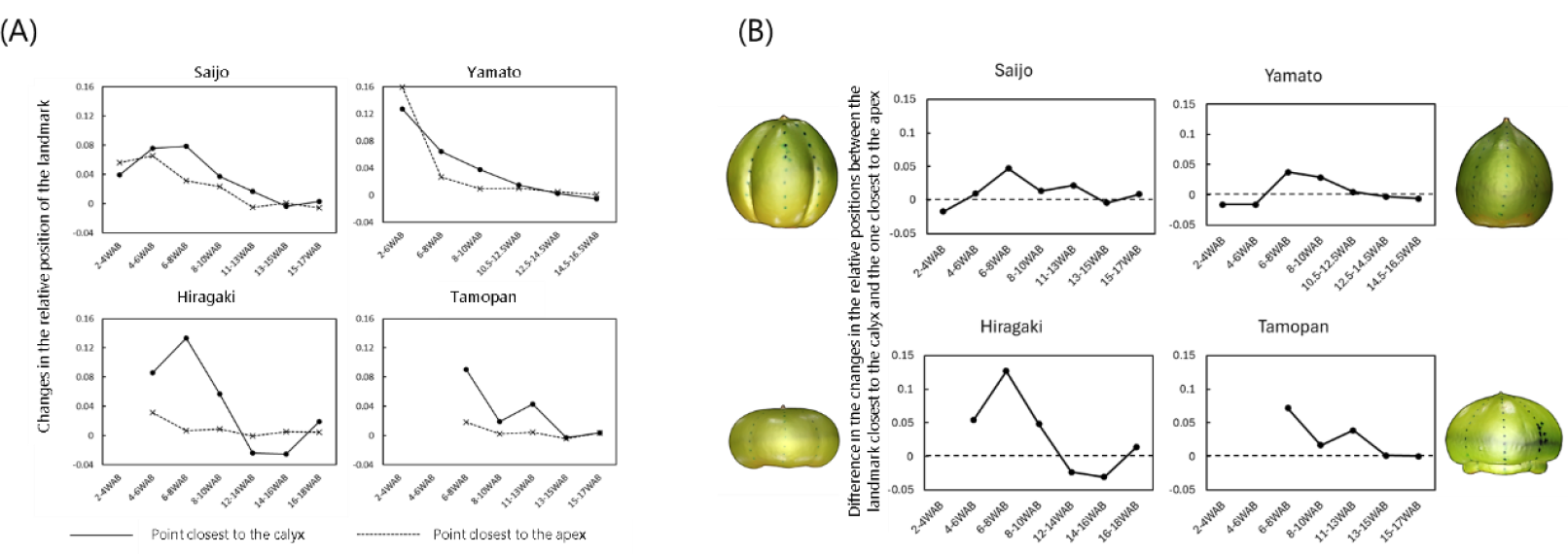
Differences in the fruit development rate around calyx and apex. (A) Changes in the relative position of the landmark closest to the calyx (solid line) and the one closest to the apex (dashed line). Those parameters were measured based on the method illustrated in Fig. 2. The surface landmark movement toward the apex was recorded as a positive change. (B) Differences in the landmark movement between the calyx and the apex ends. They were calculated by subtracting the change in the relative position of the landmark closest to the apex from that of the mark closest to the calyx.

### 3.3 Characterization of growth rates across different fruit regions in cultivars with contrasting fruit shapes

The variations in growth rates across three fruit regions: distal, middle, and proximal were summarized in Fig. 6. In all cultivars, the growth rate was highest during the early stages of development and gradually decreased thereafter in all fruit regions. By examining differences in growth rates across fruit regions, it was observed that the growth rate during the early to mid-developmental stages typically followed the order: proximal > middle > distal. This indicates a common growth gradient among fruit portions in persimmons. This gradient was more pronounced in ‘Saijo,’ ‘Yamato,’ and ‘Tamopan’; however, by approximately 16 WAB, although growth continues, this gradient became very small.

**Fig. 6.**
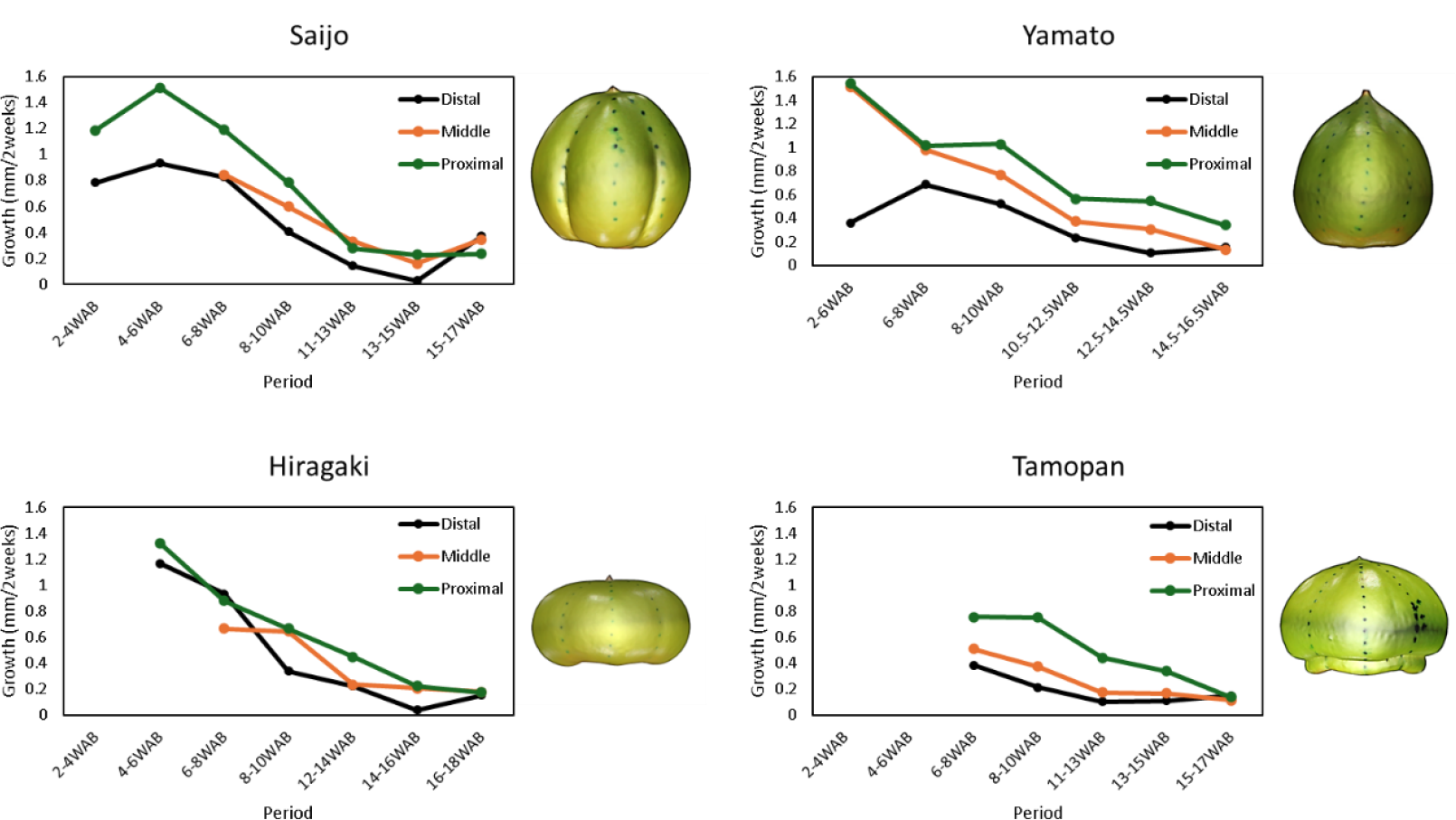
Growth rate of different fruit portions in four tested cultivars. All the surface landmarks were categorized into three, distal, middle or proximal portions depending on their positions. When there are less than three marks, they were classified only into distal and proximal portions.

Variations in the growth gradient appeared to be correlated with the aspect ratio of matured fruit. In ‘Saijo,’ the proximal portion initially exhibited the most rapid growth. However, as development progressed, its growth rate declined to a level comparable to that of the middle portion, ultimately resulting in minimal differences in growth rates among the three portions in the later stages. Conversely, in ‘Yamato,’ the middle portion exhibited a growth rate comparable to that of the proximal portion during the early developmental stages. However, along with the development, growth in the middle portion declined to a level closer to that of the distal portion, while only the proximal portion maintained noticeable growth until the later stages. A similar trend was observed in ‘Tamopan.’ In contrast, ‘Hiragaki’ did not exhibit a distinct pattern of differences in growth rates among fruit portions, as observed in the other three cultivars. Notably, during 6–8 WAB, the growth rate of the proximal portion was lower than that of the distal portion, with the middle portion showing the slowest growth. At first glance, this appears to contradict the earlier result, which indicated more vigorous growth in the proximal end than the distal end based on the analysis of landmarks closest to those ends (Fig 5). However, this discrepancy can be explained by differences in the measurement approach. The relative positional change of the landmark closest to the calyx in Fig. 5A was defined as the distance from the apex to that mark, expressed as a ratio of the total length from the apex to the shoulder of the fruit. In contrast, the tissue growth rate calculated here (Fig. 6) was based on the distance between adjacent landmarks, excluding the distance between the outermost landmark and the calyx-end. This suggests that in ‘Hiragaki’ fruit, vigorous growth occurs in a very specific and limited region between calyx and its nearest landmark in this study, but it diminishes even a short distance away from the calyx. Consequently, the average growth rate of the proximal portion remained lower, compared to the distal and middle portions.

Taking together, these findings indicate that in all cultivars, tissue growth rates were higher in the proximal end than in the distal end, suggesting that fruit development is primarily driven by growth near proximal end, in other words, calyx. The results also indicated the presence of a growth gradient, with decreasing growth rates toward the apex. However, in the extremely flattened fruit of ‘Hiragaki,’ only a specific portion around the proximal region exhibited an exceptionally high growth rate, making this gradient less apparent than in other cultivars.

## 4. Discussions

A distinctive growth gradient in persimmon fruit has long been suggested (Fujimura, 1935; Yamamura, 1984). The present study extended the previous research by utilizing 3D phenotyping of four cultivars with diverse fruit shapes, incorporating more intense data collection to allow for more precise quantification of spatial variations across time point and genotypes. The results obtained in this study are consistent with their findings, demonstrating that regardless of the cultivar, fruit tissue near the calyx end exhibited more active growth than that near the apex end, and this growth rate gradient persisted until approximately 13 WAB, which corresponds to the end of first phase of fruit growth. This also aligned with the findings of our previous study (Kusumi et al. 2024), which showed that the upper side of the horizontal groove near the calyx exhibited more rapid development during the early stages of fruit growth. These results, together with our current findings, further support that the primary site of early persimmon fruit development is located on the proximal side.

This growth pattern may have some variations among cultivars with different fruit shapes (Fig. 7). As shown in Fig. 4, ‘Hiragaki’ has an extremely flattened fruit shape, while ‘Tamopan’ exhibits a relatively flattened shape. In contrast, ‘Saijo’ and ‘Yamato’ have elongated fruit shapes (Fruit Tree Experiment Station of Hiroshima Prefecture, 1979). Considering the findings obtained in the spatial fruit growth analysis (Fig. 5 and Fig. 6), it was suggested that in flattened fruits, tissue growth was highly concentrated in a limited region near the calyx, with minimal growth extending toward the apex. On the other hand, in elongated fruits, not only was the growth most vigorous near the calyx, but growth was also relatively sustained in regions closer to the apex (Fig. 5). As a result, the growth rate gradient from the calyx to the apex was considered to be more evenly distributed in elongated fruits, while flattened fruits displayed a more pronounced gradient (Fig. 7).

**Fig. 7.**
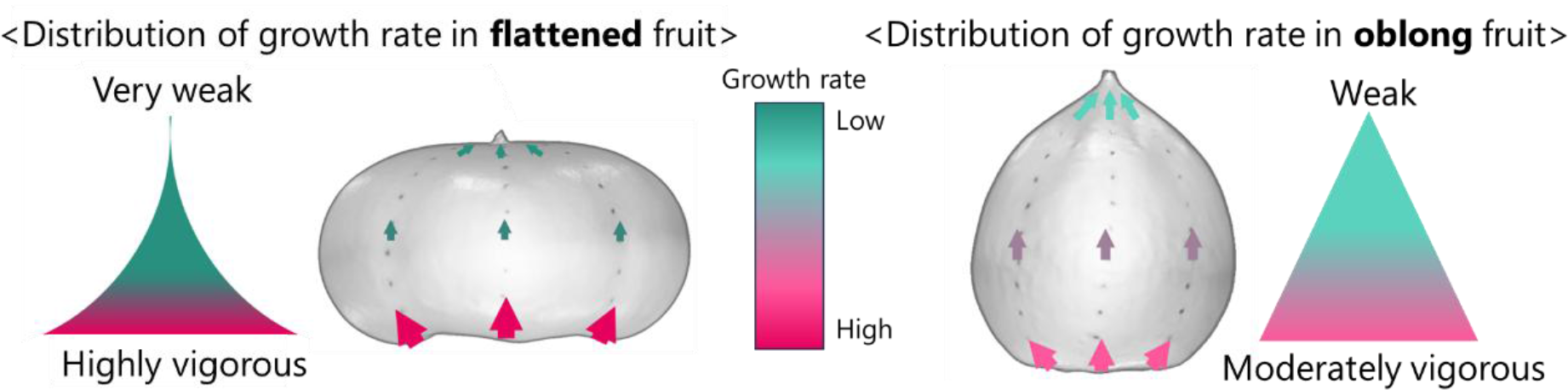
An illustration of the proposed growth gradient distribution model in flattened and oblong fruit types.

The mechanism behind the growth gradient across fruit portions has been started to reveal in *Arabidopsis*. By live-imaging the cell growth in developing *Arabidopsis* ovary, two growth gradients, horizontal and vertical, were identified (Gómez-Felipe et al., 2024). Those gradients were finely coordinated depending on the developmental stages and the identities of organs around, associated with the auxin flow. Zhang et al. (2025) also highlighted the importance of auxin flow and gradient for *Capsella* to have heart-shaped fruit. Thus, visualizing the auxin flow and gradient in developing persimmon fruit may be the next research target to understand the molecular mechanisms underlying proximal-dominant fruit development. Besides, the calyx lobes have been reported to play a critical role in persimmon fruit development, with active gas exchange during the early stage of fruit development, which is thought to enhance fruit growth by increasing metabolic activity within the fruit (Yonemori et al., 1995). This physiological function may be related to the growth rate gradient observed in this study.

Since few studies have investigated the primary sites of active development within a fruit in tree-species, it may be interesting to apply the method employed in this study to other fruit species to assess growth rate differences among fruit regions and to facilitate comparative analyses among commercially important species, such as *Prunus* and *Citrus* fruits. As these fruits lack a well-developed calyx, comparative studies may provide insights into the functional role of the calyx in fruit development. Furthermore, the application of non-destructive three-dimensional reconstruction methods to track the growth of the same fruit throughout its developmental stages would enable a more detailed modeling of tissue growth dynamics at different fruit regions over time.

Finally, along with genetic approaches taken so far, a deeper understanding of fruit growth obtained through this type of morphometric approach may have significant agronomic importance. By enabling the manipulation of tissue growth in specific portions, control of fruit size and shape may be accelerated. Moreover, it may also provide new insights into the mechanisms underlying common physiological disorders in persimmon, such as calyx-end and apical-end cracking (Yamada et al., 1987) and shallow concentric skin cracks (Iwanami et al., 2002; Yamamura et al., 1984), which are thought to result from imbalances in tissue growth rates among different fruit regions. Based on the foundation on the spatial growth variation established by this study, future investigations incorporating physiological data will be critical for applying these insights to breeding and cultivation practices.

## CRediT authorship contribution statement

**Kusumi Akane**: Investigation, Methodology, Writing – original draft. **Soichiro Nishiyama**: Conceptualization, Methodology, Resources, Supervision, Writing – review and editing. **Hisayo Yamane**: Supervision, Resources, Writing – review and editing. **Ryutaro Tao**: Conceptualization, Supervision, Resources, Writing – review and editing.

## Declaration of competing interest

The authors declare that they have no known competing financial interests or personal relationships that could have appeared to influence the work reported in this paper.

## Funding

This work was supported by the Japan Society for the Promotion of Science KAKENHI grant no. 21KK0269 to SN and 24KJ1497 to AK.

## Notes

### Competing Interest Statement

The authors have declared no competing interest.

